# Genetic Screen Identified Prmt5 as a Neuroprotection Target against Cerebral Ischemia

**DOI:** 10.1101/2023.06.26.546470

**Authors:** Haoyang Wu, Peiyuan Lv, Jinyu Wang, Brian Bennett, Jiajia Wang, Pishun Li, Yi Peng, Guang Hu, Jiaji Lin

## Abstract

Epigenetic regulators present novel opportunities for both ischemic stroke research and therapeutic interventions. While previous work has implicated that they may provide neuroprotection by potentially influencing coordinated sets of genes and pathways, most of them remains largely uncharacterized in ischemic conditions. In this study, we used the oxygen-glucose deprivation (OGD) model in the immortalized mouse hippocampal neuronal cell line HT-22 and carried out an RNAi screen on epigenetic regulators. We identified Prmt5 as a novel negative regulator of neuronal cell survival after OGD, which presented a phenotype of translocation from the cytosol to the nucleus upon oxygen and energy depletion both *in vitro* and *in vivo*. Prmt5 bound to the chromatin and a large number of promoter regions to repress downstream gene expression. Silencing Prmt5 significantly dampened the OGD-induced changes for a large-scale of genes, and gene ontology analysis showed that Prmt5-target genes were highly enriched for Hedgehog signaling. Encouraged by the above observation, we treated mice with middle cerebral artery occlusion (MCAO) with the Prmt5 inhibitor EPZ015666 and found that Prmt5 inhibition sustain protection against neuronal death *in vivo*. Together, our findings revealed a novel epigenetic mechanism of Prmt5 in cerebral ischemia and uncovered a potential target for neuroprotection.

## Introduction

Ischemic stroke, which accounts for 87% of all strokes, is the second most common cause of death and the leading cause of acquired disability in adults worldwide [1]. However, current therapy represented by rapid circulation restoration only benefits ∼5% of patients due to the narrow treatment window and small vessel blockage [1]. There is an urgent need in developing neuroprotective drugs that provide an alternative strategy to minimize neural cell loss and maximize the potential for recovery, which finally extend the therapeutic window for recanalization and improve clinical outcomes after stroke. To develop such effective therapies, genomic approaches have been employed to elucidate the mechanism of ischemic stroke-induced brain damage and the underlying molecular basis. In particular, genome-wide association, genome sequencing, transcriptomic, proteomic and metabolomic studies have identified many genomic loci, genes and pathways that are associated stroke, providing potential diagnostic biomarkers and therapeutic targets [2, 3]. Intriguingly, increasing evidence suggests that epigenetic alterations play important roles in the pathogenesis of stroke [4]. Epigenetic alterations are reversible changes in DNA or histone modifications, chromatin structure, and non-coding RNAs, and serve critical roles in the regulation of gene activity. For example, global DNA methylation, histone methylation, histone acetylation, microRNAs, enhancer RNAs, and circular RNAs were found to show significant changes after cerebral ischemia in animal models or human patients [4]. Furthermore, inhibition of DNA methyltransferases or histone deacetylases by genetic or chemical mechanisms were shown to protect against stroke induced brain damage [5]. Polycomb repressive complex genes were found to play important roles in neuroprotection after stroke [6]. Thus, epigenetic regulators present novel opportunities for both stroke research and therapeutic interventions, and they may provide neuroprotection by potentially influencing coordinated sets of genes and pathways.

Both animal and cell culture models have been developed to facilitate the study of cerebral ischemia. *In vivo*, middle cerebral artery occlusion (MCAO) in rodents can produce reliable and well-reproducible infarcts and became a commonly used approach to mimic human ischemic stroke. *In vitro*, many cell-based models, using cells treated with chemical inhibitors, enzymatic induction, or hypoxia and energy depletion, were developed to replicate key features of ischemia [7]. Even though they cannot recapitulate the complex response of stroke in intact animals, they serve as convenient and amenable tools to study ischemia-reperfusion injuries via biochemical, genetic and genomic methods. For example, oxygen-glucose deprivation (OGD) treatment has been widely used to examine the cellular mechanisms involved in cerebral ischemia and to identify potential neuroprotective agents and pathways [8]. In this study, we used the OGD model in the immortalized mouse hippocampal neuronal cell line HT-22 and carried out an RNAi screen on epigenetic regulators. We identified Prmt5 as a novel negative regulator of neuronal cell survival after OGD. We further showed that it promoted cell death by repressing genes involved in Hedgehog signaling, and its inhibition provides protection against ischemic injuries both *in vitro* and *in vivo*. Together, our findings revealed a novel epigenetic mechanism that contributes to neuronal cell death in cerebral ischemia and uncovered a potential target for neuroprotection.

## Results

### 2.1 RNAi screen identified Prmt5 as a negative regulator of neuron survival after OGD

To systematically identify epigenetic factors that regulate cellular responses after cerebral ischemia, we used the well-established OGD protocol as an *in vitro* model for ischemic injury in the mouse hippocampal neuronal cell line HT-22. We carried out an RNAi screen using a custom shRNA library targeting selected epigenetic regulators. We first optimized the OGD treatment procedure on HT-22 cells and examined the length of OGD on cell survival (Figure S1A). We chose 8 hrs of oxygen and glucose deprivation followed by 24 hrs of normal culture for the screen, because it caused significant amount of cell death with small but consistent percentage of surviving cells. Next, we selected 125 genes involved in epigenetic and chromatin regulation and generated an shRNA library targeting these genes in the pLKO.1 vector (Table S1). To perform the screen (Figure 1A), we transduced HT-22 cells with the shRNA library, selected for puromycin-resistant cells, and split the cells into two equal populations. One population was frozen as a pellet and used later as the library-transduced cells. The other was treated with the optimized OGD procedure. After the treatment, the cells were cultured under normal conditions and expanded for another 10 days to allow those that survived OGD to grow. The OGD-treated cells were collected and frozen as a pellet as well. We extracted genomic DNA from both the library-transduced and the OGD-treated cell pellets, polymerase chain reaction (PCR)-amplified the shRNAs from the genomic DNAs, and sequenced the shRNA pools from the two cell populations using the Illumina platform. Based on the representation of the shRNAs in the cells, we found that most of the shRNAs were depleted after OGD treatment, suggesting that the genes they target are likely required for cell survival during or after OGD. Importantly, we also found a small number of shRNAs that showed increased representations after OGD, suggesting that the inhibition of their target genes likely promoted cell survival. Encouragingly, shRNAs against Dnmt3b, Pcgf6 and Mga were among those that are enriched after OGD treatment (Figure 1B), consistent with the notion that DNA methylation and polycomb repressive complexes play important roles in stroke injury and neuroprotection [9-12].

**Figure 1.**
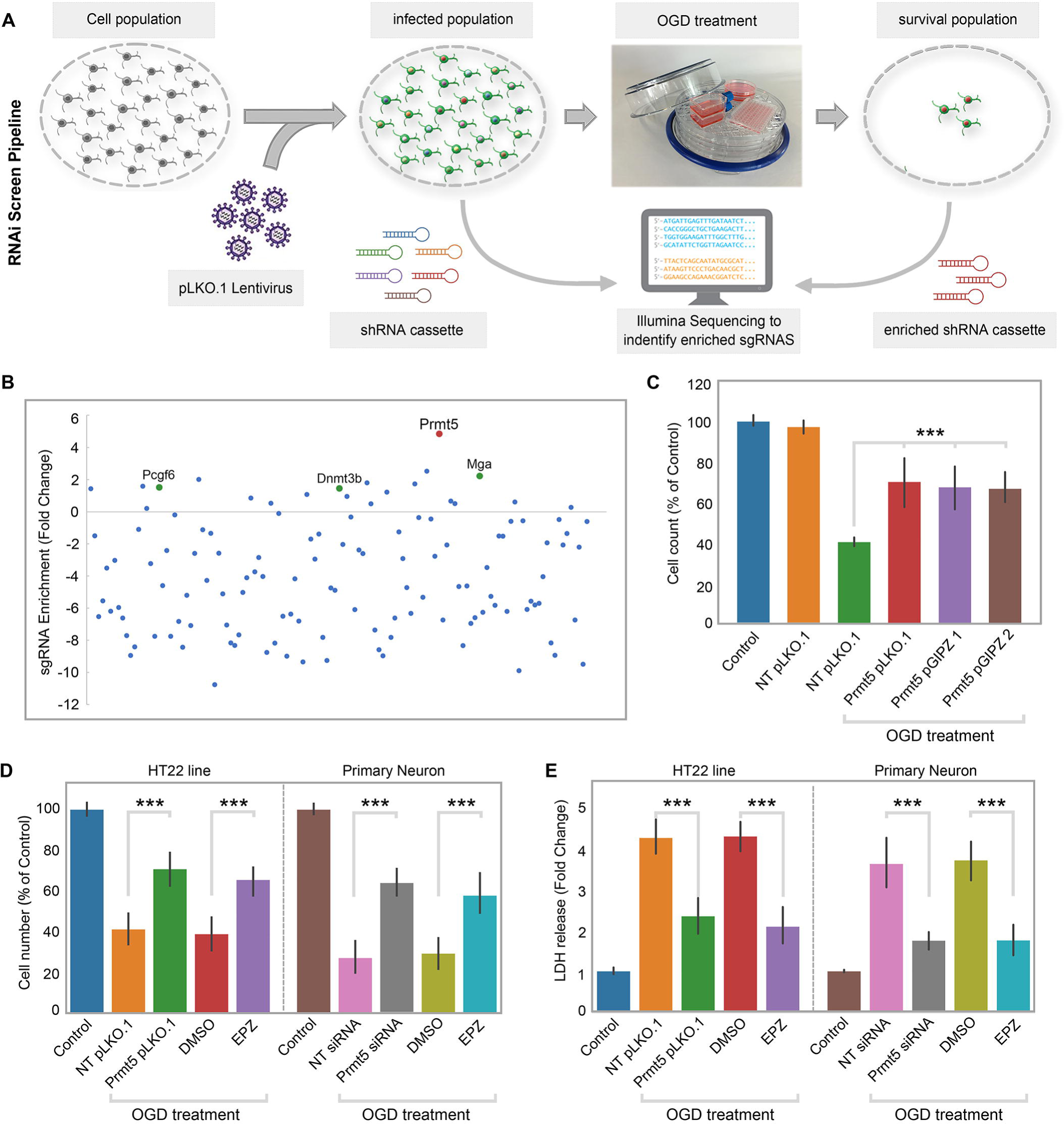
RNAi screen identified Prmt5 as a negative regulator of neuron survival after OGD. **(A)** Overview of the study methodology. **(B)** shRNA representation after OGD treatment. Prmt5 sgRNA have the highest enrichment after OGD treatment. **(C)** Effect of Prmt5 silencing in HT-22 cells after OGD. HT-22 cells (Control) or HT-22 cells transduced with the indicated shRNA lentiviruses were cultured with or without OGD treatment. Cell numbers were counted after the treatment and normalized against the control. Compared with non-targeting shRNA lentiviruses (NT pLKO.1), Prmt5 knockdown promoted a significant cell survival after OGD. **(D-E)** Effect of Prmt5 silencing of inhibition in HT-22 cells and primary neurons after OGD. Cell viability was determined using the Cell Counting Kit-8. Cellular cytotoxicity was determined using the Lactate Dehydrogenase Activity Assay Kit. Compared with NT pLKO.1, Prmt5 pLKO.1 promoted a significant cell viability. Similar result could be found in Prmt5 inhibitor EPZ015666 (EPZ) *vs.* solvent control (DMSO). Results were plotted as mean ± SEM. \*\*\**p* < 0.001.

Among the enriched shRNAs, the shRNA against Prmt5 displayed the largest fold-change (Figure 1B). Therefore, we decided to focus on Prmt5 and set out to validate its role in ischemic cell death. To rule out potential off-target effects by the Prmt5 shRNA in the screen, we designed additional shRNAs against Prmt5 in a different vector pGIPZ. We found that Prmt5 knockdown (KD) by the original pLKO.1 shRNA as well as the two new pGIPZ shRNAs all led to significant improvement in HT-22 cell survival after OGD (Figure 1C). Moreover, treatment of the cells with a small molecule Prmt5 inhibitor EPZ015666 resulted in similar outcomes (Figure 1D), further supporting the fact that Prmt5 is responsible for the protective effect. Consistent with the above results based on cell numbers, Prmt5 inhibition significantly reduced OGD-induced HT-22 cell death in the lactate dehydrogenase (LDH) cytotoxicity assay (Figure 1E). Finally, in addition to the immortalized HT-22 cell line, Prmt5 shRNA or inhibitor promoted the survival of primary mouse neurons in culture after OGD treatment (Figure 1D-F). Together, these results clearly demonstrated that Prmt5 plays an important role during OGD and its inhibition protects neuronal cells against OGD-induced damage.

### 2.2 Prmt5 protein translocates into the nucleus upon ischemic injury

To understand the molecular function of Prmt5 in OGD, we examined the behavior of its protein product by imaging in HT-22 cells. We found that PRMT5 is predominantly localized in the cytoplasm under normal conditions. Upon OGD treatment, however, it relocates to the nucleus and becomes enriched in the nuclear compartment (Figure 2A). To rule out any potential artifacts caused by the antibody, we knocked-in an HA-tag to the C-terminus of the endogenous Prmt5 gene in HT-22 cells using CRISPR-mediated genome editing, and re-examined Prmt5 localization with the HA antibody. Similarly, we found that the endogenous HA-tagged Prmt5 translocates from the cytosol to the nucleus after OGD (Figure 2A). To complement the imaging results, we carried out biochemical fractionation of the HT-22 cells and separated the cellular content into cytoplasmic, nucleoplasmic and chromatin-bound fractions. Consistent with the above, Prmt5 protein moved from the cytoplasm to the nucleus in OGD-treated cells (Figure 2B). Further, we found that Prmt5 protein became largely chromatin-bound, suggesting that it may participate in OGD-induced transcriptional responses. Finally, we repeated the experiments in OGD-treated primary mouse neurons in culture and observed the same result (Figure 2CD).

**Figure 2.**
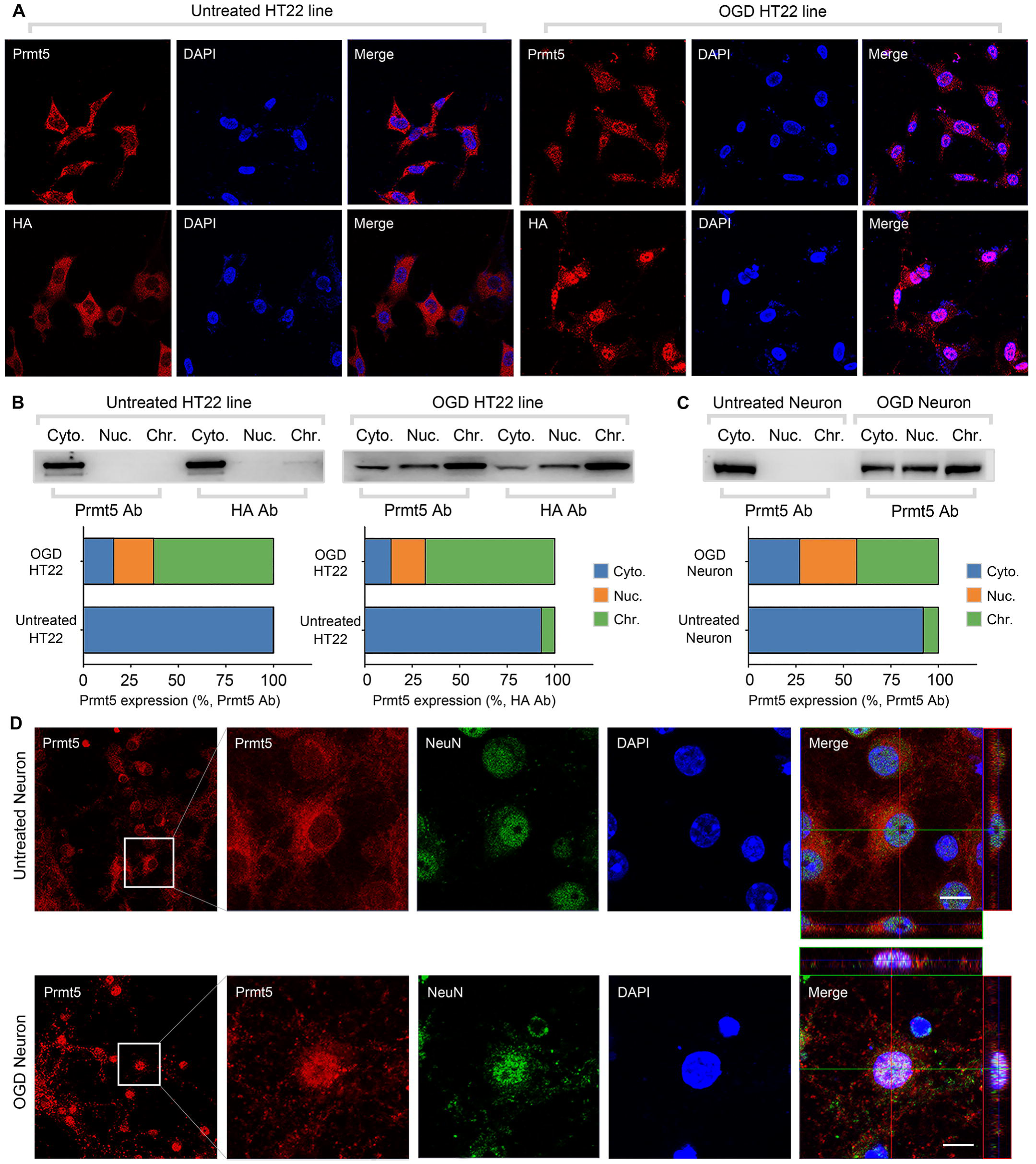
Oxygen-glucose deprivation led to Prmt5 nuclear translocation *in vitro*. **(A)** Immunofluorescence staining to show Prmt5 cellular localization after OGD in HT-22 cells. HT-22 cells were stained with the Prmt5 antibody or HA antibody for C-terminal HA-tagging of the endogenous Prmt5 by CRIPR-mediated genome editing. Cell nuclei were counterstained by DAPI. **(B)** Biochemical fractionation to show Prmt5 cellular localization after OGD. Prmt5 in each fraction was detected by Western blot, using either the Prmt5 or HA antibody. **(C-D)** Primary cortical neurons were treated with or without OGD and subjected to biochemical fractionation. Primary neurons were stained with the Prmt5 or NeuN (neuronal marker) antibody. Cyto., Nuc. and Chr. stand for cytoplasmic, nucleoplasmic and chromatin-bound fractions

Next, we wanted to test whether Prmt5 translocation also happens *in vivo*. We used the left middle cerebral artery occlusion (MCAO) model to mimic ischemic injury in the mouse brain, and examined Prmt5 localization in both the ipsilateral and contralateral side by immunohistochemistry. Similar to what was observed in cultured cells, Prmt5 signal was mostly detected in the cytoplasm on the contralateral side, but became highly enriched in the nucleus on the ipsilateral side (Figure 3A). Upon closer examination, Prmt5 nuclear translocation was detected in both the penumbra area and ischemic core of the ipsilateral side (Figure 3B). We further verified these observations by immunofluorescence staining (Figure 3C). Importantly, biochemical fractionation of the neuronal tissues again showed that Prmt5 not only translocates into the nucleus but also binds to the chromatin upon ischemic injury (Figure 3D). As Prmt5 has been implicated in chromatin and gene regulation, these results strongly suggested that Prmt5 may enter the nucleus and bind the chromatin to regulate neuronal gene expression upon OGD.

**Figure 3.**
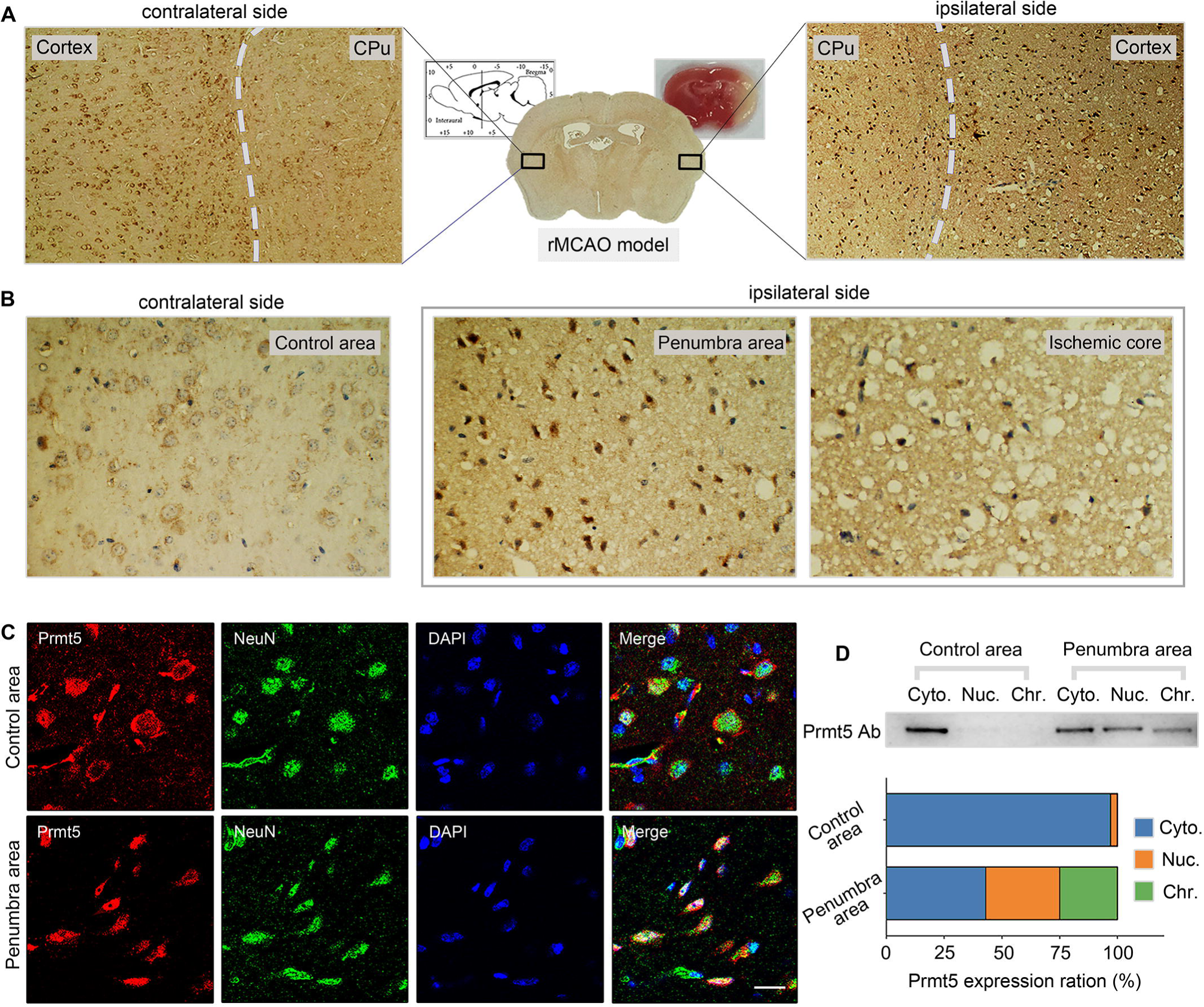
Middle cerebral artery occlusion led to Prmt5 nuclear translocation *in vivo*. **(A-C)** Immunohistochemical and Immunofluorescence staining to show Prmt5 cellular localization in ipsilateral and contralateral sides after MCAO. **(D)** Biochemical fractionation to show Prmt5 cellular localization after MCAO. Prmt5 in each fraction was detected by Western blot, using either the Prmt5 antibody. Cyto., Nuc. and Chr. stand for cytoplasmic, nucleoplasmic and chromatin-bound fractions

### 2.3 Prmt5 represses Hedgehog signaling expression after OGD

To test whether and how Prmt5 regulates gene expression after OGD, we carried out RNA-seq in OGD-treated HT-22 cells *vs.* control, as well as OGD-treated HT-22 cells with Prmt5 KD *vs.* control. We found that OGD treatment led to the up-regulation of 2,186 genes and down-regulation of 2,926 genes (Figure 4A, Table S2). Interestingly, while Prmt5 KD alone had minimal impact on gene expression, it significantly dampened the OGD-induced changes for a large number of genes (Figure 4B, Table S2). Specifically, 1,035 of the 2,186 OGD-induced genes and 2,090 of the 2,926 OGD-repressed genes were reversed or partially reversed upon Prmt5 KD (Figure 4B, Table S2). This is consistent with the cellular phenotype, in which Prmt5 KD partially rescued OGD-induced cell death.

**Figure 4.**
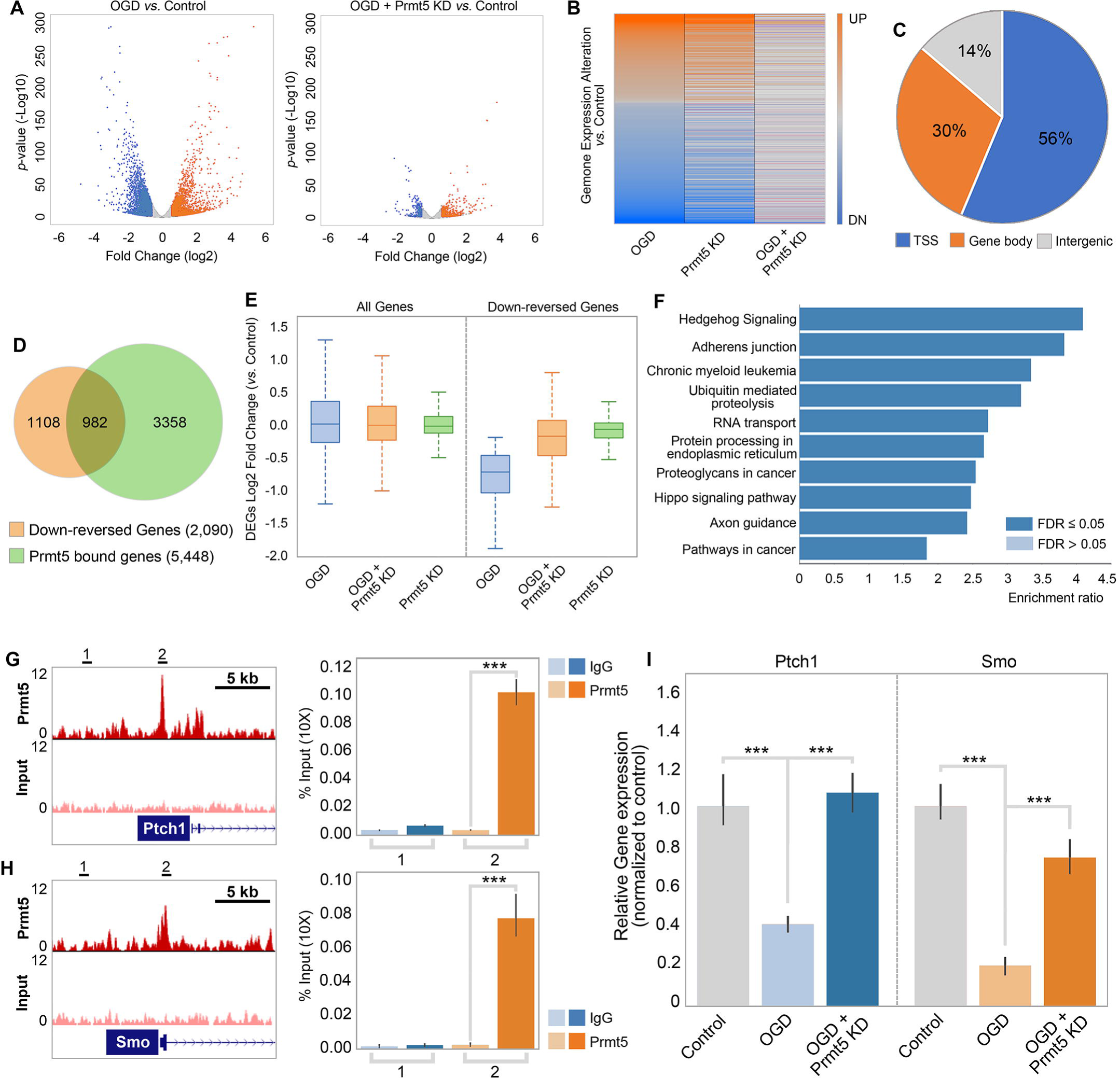
Prmt5 represses Hedgehog signaling expression after OGD. **(A)** Gene expression changes after OGD. HT-22 cells were transduced with Control (non-targeting) or Prmt5 shRNA lentiviruses, drug selected, and treated with OGD. Cells were collected and total RNAs were extracted for RNA-seq. Differentially expressed genes (DEGs) after OGD in the control cells were identified by log2-fold change >1 and *p*-value < 0.01. Orange: up-regulated genes after OGD; Blue: down-regulated genes after OGD. **(B)** Effect of Prmt5-KD on OGD-induced gene expression changes. OGD-induced DEGs were defined as described in (A), and the expression changes of the DEGs after OGD in control and Prmt5-KD cells were plotted by heat map. **(C-D)** Genomic regions occupied by Prmt5 based on ChIP-seq in HT-22 cells treated with OGD. Reversal of OGD-induced gene expression changes by Prmt5-KD. Prmt5-target genes were defined as described in the text. **(E)** Expression changes of all detected genes or Prmt5-target genes after OGD in control or Prmt5-KD cells were examined by box plots. **(F)** Kyoto Encyclopedia of Genes and Genomes (KEGG) pathway analysis of Prmt5-target genes. **(G-H)** Genome browser tracks of Prmt5 occupancy at selected Prmt5-target genes. ChIP-qPCR of Prmt5 occupancy in HT-22 cells treated with OGD at selected Prmt5-target genes (Ptch1 and Smo). **(I)** RT-qPCR to show the expression of selected Prmt5-target genes in control or Prmt5-KD HT-22 cells upon OGD treatment. \*\*\**p* < 0.001.

As shown in Figure 2, OGD treatment resulted in Prmt5 nuclear translocation and chromatin binding. To test whether Prmt5 directly regulates gene expression and contributes to OGD-induced transcriptional changes, we carried out Prmt5 chromatin immunoprecipitation followed by high throughput sequencing (ChIP-seq) in HT-22 cells treated with OGD. In total, we identified 16,061 peaks bound by Prmt5. Interestingly, a significant fraction of the Prmt5-bound genomic regions (56%) are near the transcription start sites (TSSs) (Figure 4C, Table S2), consistent with the notion that Prmt5 may regulate downstream gene expression. Indeed, when intercepted with the RNA-seq data, we found that Prmt5 occupies the promoter regions of a large fraction of the 2,090 genes that were down-reversed genes in OGD (Figure 4D, Table S2). Furthermore, the OGD-repressed genes appeared to be dependent on Prmt5, as Prmt5 KD largely reversed their expression changes in OGD (Figure 4E). Therefore, we defined these genes that are bound by Prmt5, down-regulated after OGD treatment but rescued by Prmt5 KD as Prmt5-target genes in OGD. And the above results suggested that under OGD conditions, Prmt5 enters the nucleus and suppresses the expression of its target genes. Gene ontology analysis showed that Prmt5-target genes are highly enriched for Hedgehog signaling (Figure 4F). To validate the above findings based on genomic data, we carried ChIP quantitative PCRs (ChIP-qPCRs) and reverse transcription quantitative PCRs (RT-qPCRs). on selected OGD-repressed Prmt5-target genes, including Ptch1 and Smo. We found that Prmt5 indeed occupied their promoter regions (Figure 4GH) and is responsible for their down-regulation after OGD treatment (Figure 4I). Together, our findings strongly supported the notion that upon oxygen and glucose deprivation, Prmt5 promotes neuronal cell death by repressing the expression of genes involved in hedgehog signaling. Prmt5 silencing provides neuroprotection by preventing the inhibition of hedgehog pathways.

### 2.4 Prmt5 inhibition promotes neuron survival after ischemic injury in mouse brain

To test the relevance of our model *in vivo*, we first examined the behavior of the Prmt5-target genes in OGD we identified in ischemic injury in mice. We used a public RNA-seq dataset generated from a MCAO mouse model and carried out gene set enrichment analysis (GSEA). We found that Prmt5-target genes in OGD are highly enriched for those that were down-regulated after cerebral infarction in the MCAO mouse (Figure 5A), suggesting that our model may apply in animals.

**Figure 5.**
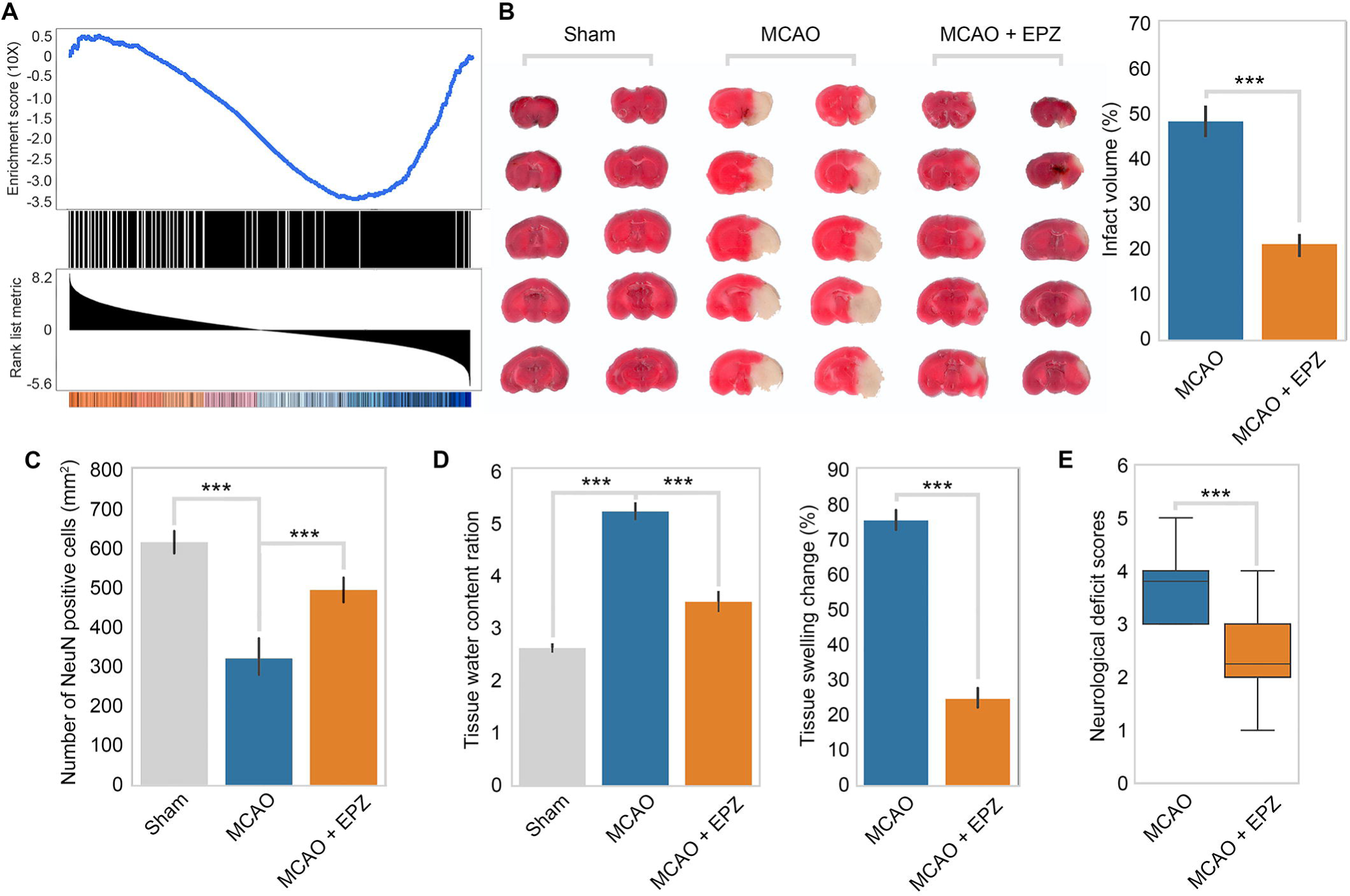
Prmt5 inhibition promotes neuron survival after ischemic injury in mouse brain. **(A)** Gene set enrichment analysis for Prmt5-target genes in the MCAO mouse stroke model. Gene expression changes after MCAO was based on 29862458. **(B)** Effect of Prmt5 inhibition on infarct volume in the MCAO model. Mice were treated with subjected with sham (Sham group), MCAO with solvent (MCAO group) or MCAO with the Prmt5 inhibitor EPZ015666 (MCAO + EPZ group). The infarct volume was determined by TTC staining. **(C-E)** For tissue immunofluorescence staining, samples were fixed using 4% paraformaldehyde at room temperature for 15 min, followed by 0.5% Triton X-100 permeabilization for 10 min and 0.5% bovine serum albumin blocking for 30 min. The mice were sacrificed for NeuN immunofluorescence staining, and positive staining cell were count as survival neurons. Tissue water content ration, tissue swelling change and neurological deficit score were determined as described in the methods. \*\*\**p* < 0.001.

Encouraged by the above observation, we set out to test whether Prmt5 inhibition may provide protection against neuronal cell death in mouse brain after ischemic injury as it did in cultured cells. We treated adult mice with either solvent only or the Prmt5 inhibitor EPZ015666 intranasally and then induced ischemic injury in the left side of the brain with the MCAO procedure. After the surgery, we first assessed the neurological damage based on behavioral changes and then sacrificed the animals and examined the damages in their brain tissue. Indeed, we found that Prmt5 inhibition resulted in significant protective effects. Specifically, EPZ015666 treatment reduced the infarct volume (Figure 5B) and increased the number of NeuN-positive neurons after MCAO (Figure 5C). It also reduced the water content and tissue swelling caused by infarction (Figure 5D). Most importantly, we found that EPZ015666 treatment led to significantly improved functional recovery in the animals as evidenced by the reduction in the neurological deficit scores (Figure 5E). Therefore, these results strongly suggested that Prmt5 plays a critical role in mediating neuronal cell death after ischemic injury in the brain and may serve as a novel therapeutic target for stroke treatment.

## Discussion

While genome-wide association and expression profiling studies have uncovered many genomic loci, genes, and regulatory elements involved in stroke, the causal factors underlying stroke-induced damage are often confounded by the large number of incidental events accompanying stroke. Therefore, new approaches to define genes with functional relevance are essential to understand stroke and identify potential therapeutic targets. In that sense, genetic screens can provide a more direct strategy to search for genes that support neuronal cell survival during stroke conditions [13]. The development of RNAi and CRIPSR technologies, as well as the multiplex screening method, further promoted fast and convenient screens in mammalian cells [14]. Here, we carried out a small-scale RNAi screen in HT-22 cells and identified a list of epigenetic factors that regulate neuronal cell death under ischemic conditions. Our study illustrates the power of forward genetics for the systematic study of ischemic stroke-induced injuries and presents a new strategy for the identification of neuroprotection agents.

Among the hits from the screen, we focused on Prmt5, as its silencing presented the strongest phenotype. Prmt5 belongs to the evolutionarily conserved arginine methyltransferase protein family [15]. It catalyzes the transfer of a methyl group from S-adenosyl-methionine to the guanidino nitrogen atoms of arginine as monomethylation and symmetric demethylation [15]. Prmt5 has been implicated in various developmental processes and diseases [16]. In particular, it is overexpressed in many cancers and elevated Prmt5 expression often correlates with poor prognosis [17]. Conversely, its depletion or inhibition leads to reduced proliferation and increased apoptosis in cancer cells [17]. In addition, Prmt5 is also required for early embryonic development and the maintenance of embryonic and neural stem cells [18]. However, we found that Prmt5 silencing by RNAi or inhibition by a small molecule enhances neuronal cell survival upon ischemic injury both *in vitro* and *in vivo*. Consistent with our findings, Prmt5 silencing was reported to reduce oxidative stress-induced cell death in renal ischemia [19]. Thus, Prmt5 appears to play dual roles in cell viability in a context-dependent fashion, being pro-proliferation in cancer and stem cells but pro-cell death under ischemic conditions.

Mechanistically, we showed that Prmt5 protein translocates from the cytosol to the nucleus upon oxygen and energy depletion both *in vitro* and *in vivo*. Like other arginine methyltransferase family members, subcellular localization of Prmt5 protein appears to be cell type specific [20]. In embryonic stem cells, it is predominantly cytoplasmic in self-renewing condition, but relocates to the nucleus upon differentiation [21]. During mammalian embryonic development, it shuttles between the cytosol and the nucleus to fulfill its many functions in gene regulation and epigenome silencing [22, 23]. In cancer, Prmt5 protein was found to show different cellular localization between normal and tumor tissues and between tumor types [24, 25], suggesting that its compartment-specific functions may regulate distinct molecular programs. In our study, we showed that Prmt5 is highly expressed in neurons and localizes predominantly in the cytoplasm under normal conditions, consistent with previous reports [26, 27]. Under OGD or ischemic conditions, it relocates to the nucleus and contributes to neuronal cell death. As transcription factors can interact with Prmt5 and regulate its cellular localization [28], we speculate that Prmt5 may be recruited to the nucleus to carry out its function in transcriptional regulation by factor(s) in response to ischemia.

Prmt5 plays important roles in gene regulation by methylating histones, transcription factors, and spliceosome proteins [29]. It also interacts with ATP-dependent chromatin remodelers to control gene expression [30]. We found that as Prmt5 translocates to the nucleus upon OGD, it binds to the chromatin and a large number of promoter regions to repress downstream gene expression. This is consistent with the notion that Prmt5 often acts as a transcription repressor [31]. Interestingly, OGD treatment alone caused similar numbers of genes to be up- or down-regulated. However, Prmt5-target genes are mostly down-regulated after OGD and are enriched for those involved in cell survival. Thus, OGD or stroke-induced neuronal cell death may largely be caused by the repression of survival pathways and re-activating these pathways may thereby provide an important mechanism for neuroprotection.

Hedgehog signaling is an important signaling pathways that is frequently used during development for intercellular communication, as well as regeneration and homeostasis [32]. In nervous system specific hedgehog, sonic hedgehog (Shh), canonically binds and inactivates Ptch1 inhibition of Smo for downstream Gli-1 signaling cascade, which has particularly marked roles in nervous system cell type specification and limbs patterning [33, 34]. Many studies have emphasised its impact on neural cell survival and tissue regeneration/repair after ischemic stroke. Chechneva et al. found that Shh signaling activation has promoted on neural cell survival and tissue regeneration/repair after ischemic stroke [35]. Moreover, Shh agonist could also induce a decrease in blood-brain barrier permeability and an increased of newly generated neurons and neovascularization in the MCAO model [35]. In vitro studies, the expression of of Shh, Ptch1, and Gli-1 was significantly downregulated within 24 hours after OGD exposure, which were closely associated with increasing numbers of apoptotic cells [36]. Activation of Shh signals promoted CREB and Akt phosphorylation, upregulated the expressions of BDNF, neuroligin, and neurexin, and decreased NF-κB phosphorylation following OGD [36]. Therefore, the regulation of Prmt5 on Shh signaling may be one of the important mechanisms of its ischemia protection.

As mentioned above, Prmt5 plays critical roles in tumorigenesis and is therefore a therapeutic target that is actively pursued for cancer treatment. Several Prmt5 inhibitors, some of which highly potent, are in clinical trials to test their safety and efficacy [37]. It will be interesting to systematically test their effect in protecting against neuronal cell death in cell-based and animal models of stroke, in order to gain more insights to the molecular events after stroke as well as develop potential therapeutic interventions.

There were two major limitations should be contemplated when interpreting these results. On one hand, HT-22 is an immortalized mouse hippocampal cell line, which is a sub-line derived from parent HT4 cells that were originally immortalized from cultures of primary mouse hippocampal neurons [38]. There exists to a certain extent difference from primary neurons under OGD stress and may be affected by cell proliferation regulatory elements in HT-22. On the other hand, the Prmt5 function seems to closely link to its subcellular localization. The nuclear Prmt5 plays specific roles in transcription regulation by directly modulating the activity of several transcription factors or by methylating histones (H4R3, H2AR3, H3R8, and H3R2), while the cytoplasmic expression of Prmt5 is involved in splicing, translation and regulation of receptor-ligand signaling and organelle integrity [20]. The mechanism of how prmt5 is involved in neuroprotection needs further research.

## Materials and methods

### 4.1 HT-22 cells and primary cortical neurons culture

HT-22 cells were kindly provided by Dr. Richard Dargusch (Salk). Its identity has been authenticated with Short tandem repeat (STR) profiling, and no mycoplasma or other microbiological contamination was found. Cells were routinely cultured in Dulbecco’s Modified Eagle’s Medium (Gibco) supplemented with 10% fetal bovine serum (Gibco), and the cultures were maintained at 37°C in a humidified atmosphere containing 5% CO_2_. For CRISPR-mediated gene targeting, pX330 and homologous recombination (HR) donor plasmids were co-transfected into HT-22 for HA-tag knock-in. Transfected cells were seeded at colonal density, and individual colonies were picked and screened by PCR. Correctly targeted clones were amplified and re-screened to confirm genotype. Primers and oligos used in this study were listed in Table S3. The neonatal mouse neurons were isolated as previously described with modifications [34]. Briefly, cerebral cortices were removed from 1-3 day old C57BL/6 mouse pups, stripped of meninges and blood vessels, and minced. Tissues were dissociated with 0.25% trypsin for 15 min at 37 and gentle trituration. Neurons were resuspended in a complete culture medium (Neurobasal medium containing 2% B27 supplement and 0.5 mM L-glutamine) and plated at a density of 3×10^5^ cells/cm^2^. Neurons were maintained at 37 in a humidified 5% CO_2_ incubator and half of the culture medium was changed every other day. The cultured neurons were used for experiments within 8-10 days. All experimental protocols and animal handling procedures were performed in accordance with the National Institutes of Health (NIH) guidelines for the use of experimental animals and were approved by the Animal Care and Use Committee of The Second Affiliated Hospital of Air Force Medical University.

### 4.2 OGD treatment

OGD treatment was carried out as previously described [39]. Briefly, cells were rinsed and cultured in the glucose-free Earle’s balanced salt solution (BSS) with the following composition (in mmol/L): 116 NaCl, 5.4 KCl, 0.8 MgSO_4_·7H_2_O, 1 NaH_2_PO_4_·2H_2_O, 26.2 NaHCO_3_, 0.01 glycine, 1.8 CaCl_2_· 2H_2_O, and pH 7.4. The culture was then placed in an Hypoxia Incubator Chamber (StemCell Technology) and were equilibrated for 15 minutes with a continuous flux of 95% N_2_/5% CO_2_ gas. The chamber was then sealed and placed into a humidified incubator at 37°C for 8 hours (HT-22) or 60 minutes (primary cortical neuron). OGD was terminated by removing the cultures from the chamber, replacing BSS with normal culture medium, and returning cells to the normoxic conditions for 24 hours. Cells cultured under normal conditions during the experimental period were used as controls.

For Prmt5 inhibitor treatment, EPZ015666 (Selleckchem Company) was prepared in dimethyl sulfoxide (DMSO) at a concentration of 100 mM and stored at −20 °C before use. EPZ (10μM final concentration) or DMSO was added to cells 20 minutes before the initiation of OGD and to the BSS during the OGD treatment. For siRNA transfection of the primary neurons, the culture medium was refreshed at day 7. Cells were transfected with siRNAs using Lipofectamine 3000 according to manufacturer’s instructions. The medium was replaced the next day, and OGD experiments were performed 48 h after transfection.

### 4.3 shRNA library and RNAi screen for OGD

Oligos used in this study were listed in Table S3. For pLKO.1 shRNAs, complementary single-stranded oligos were annealed and cloned as suggested by the RNAi consortium. pGIPZ shRNAs were ordered from Dharmacon. All plasmids were confirmed by sequencing. The shRNA library used for the screen was as previous described [40]. shRNA lentivirus was prepared by transfecting 293T cells using TransIT-293 (Mirus) using a standard protocol from Addgene.

For the shRNA screen, HT-22 cells were transduced with shRNA lentivirus and selected for viral infection with 2 μg/ml puromycin for 4 days. After selection, cells were split into 2 halves. One half was harvested and frozen as a pellet (Untreated). The other half was subjected to OGD treatment, replated, cultured until confluent and harvested (Treated). Genomic DNA was extracted from the cell pellets and the integrated shRNAs were amplified according to the pooled screen protocol from the Broad Institute (https://portals.broadinstitute.org/gpp/public/resources/protocols). The amplified shRNAs were sequenced on the Illumina platform, and the representations of the shRNAs in the Untreated and Treated cells were calculated to identify shRNAs that were enriched after treatment.

### 4.4 RT-qPCR and RNA-seq

Total RNA was isolated from cells using the GeneJet RNA purification kit (Thermo Scientific), and 0.5 μg total RNA was reverse transcribed to generate cDNA using the iScriptTM cDNA Synthesis Kit (Bio-Rad) according to manufacturer’s instructions. RT-qPCRs were performed using the SsoFastTM EvaGreen Supermix (Bio-Rad) on the Bio-Rad CFX-384 Real-Time PCR System. Actin was used for normalization. All experiments were performed at least three times, and representative results were shown in the figures. For RNA-seq, libraries were prepared from two biological replicates using the TruSeq RNA Sample Prep Kit and sequenced on the NextSeq (Illumina).

### 4.5 Cell viability and LDH assay

Cell viability assay was performed using the cell proliferation reagent Cell Counting Kit-8 (CCK-8) following the manufacturer’s protocol (Sigma). Briefly, cells were culture din 96-well plates and subjected to different treatments as described in each experiment. 10 μL CCK-8 was added to each well, and the plates were incubated at 37°C for 90 min. The absorbance was determined with a Microplate Reader (Bio-Rad). Neuronal cytotoxicity was determined with the Lactate Dehydrogenase Activity Assay Kit according to the manufacturer’s instructions (Sigma). Briefly, cells were culture din 96-well plates and subjected to different treatments as described in each experiment. 50 μl of supernatant from each well was collected to assay the LDH activity.

### 4.6 Subcellular fractionation and Western blot

Subcellular fractionation was adapted from Mendez and Stillman [41]. Briefly, harvested cells were washed with phosphate-buffered saline (PBS) and re-suspended in cytosolic buffer (10 mM HEPES [pH 7.4], 10 mM KCl, 1.5 mM MgCl_2_, 0.34 M sucrose, 10% glycerol, 1 mM Dithiothreitol (DTT)) supplemented with protease inhibitors at a concentration of 20-40 million cells/ml, and incubated on ice for 5 min. A total of 1% Triton-X 100 in equal volume of cytosolic buffer was added to a final concentration of 0.1%, and the cells were mixed by gently pipetting and further incubated for 10 min on ice. 10% of the cell suspension was taken as the whole-cell extract fraction. The remaining cell suspension was centrifuged at 1300 × *g* for 5 min at 4°C to separate the cell nuclei, and supernatant containing the cytoplasmic fraction was collected. Nuclei were washed once in cytosolic buffer, then lysed 10 min on ice in 1× volume chromatin extraction buffer (3 mM EDTA, 0.2 mM EGTA, 1 mM DTT) supplemented with protease inhibitors. Insoluble chromatin was pelleted by centrifugation at 1700 × *g* for 5 min at 4°C, and supernatant containing the nucleoplasm fraction was collected. The chromatin pellet was washed once with chromatin extraction buffer. All fractions were boiled in LDS loading buffer. All experiments were performed three or more times, and representative results were shown in the figures.

Cell lysates or extractions were loaded into a NuPAGE Bis-Tris gels (4–12%) and transferred onto PVDF or NC membranes. The membrane was blocked with 5% non-fat milk, followed by incubation with primary antibodies (Table S4) and HRP-conjugated secondary antibodies. Chemiluminescence signal was generated with ECL reagents (GE) and detected using ChemiDoc Touch Imaging System (Bio-Rad).

### 4.7 Immunostaining

For cell immunofluorescence staining, samples were fixed using 4% paraformaldehyde at room temperature for 15 min, followed by 0.5% Triton X-100 permeabilization for 10 min and 0.5% bovine serum albumin blocking for 30 min. Samples were then incubated with primary antibodies (Table S4) at 37°C for 2 h or 4°C overnight, followed by fluorescent secondary antibodies (Life Technologies). Nuclei were counterstained with DAPI (Vector Laboratories). For tissue immunohistochemical staining, mice were sacrificed and intracardially perfused with 4% paraformaldehyde for 10 min. Mouse brain was removed and fixed overnight at 4°C, and dehydrated in a graded ethanol series and processed for paraffin embedding. For immunofluorescence, sections were incubated with the primary antibodies overnight at 4°C, and then fluorescent secondary antibodies with DAPI nuclear staining. The specificity of the staining was confirmed by omitting the primary antibodies. For DAB staining, the images of immunohistochemical results were obtained by a DMR-X microscope coupled with a DC500 digital camera (Leica) and the image analysis system Quantimet Q550 (Leica). For immunofluorescent double labelled staining, the images were collected by a Leica SP5 confocal microscope (Leica) and recorded sequentially using Leica Application Suite Software (Leica). All experiments were performed three or more times, and representative results were shown in the figures.

### 4.8 *In vivo* model of brain ischemia and damage evaluation

This study was performed in strict accordance with the recommendations in the Guide for the Care and Use of Laboratory Animals of the National Institutes of Health. All of the animals were handled according to approved institutional animal care and use committee protocols of Air Force Medical University. The protocol was Committee on the Ethics of Animal Experiments of Air Force Medical University. All surgery was performed under sodium pentobarbital anesthesia, and every effort was made to minimize suffering. 8-12 week old mice (25-30 g) were obtained from the Laboratory Animal Center of the Air Force Medical University. MCAO was used to induce focal cerebral ischemia. Briefly, mice were anesthetized using 5% isoflurane in 30% O_2_ / 70% N_2_O and placed in the supine position on a heating pad. Ischemia was produced by advancing the tip of a rounded nylon suture into the left internal carotid artery through the left external carotid artery. After 60 min of occlusion, the thread was withdrawn to allow reperfusion. Mice were sutured and placed in a 35 °C nursing box to recover from anesthesia, and then returned to the cage. Sham group mice received midline neck incisions. The left common carotid artery was isolated, but not cut. In MCAO + EPZ groups, mice were intranasally treated with 20 mg/kg EPZ015666 1 h before surgery. The MCAO group mice were intranasally administered with an equal volume of the DMSO solution used to dissolve EPZ. After the animal was awakened, neurological damage was assessed as follows [42]: 0, no deficit; 1, forelimb weakness and torso turning to the ipsilateral side when held by tail; 2, circling to the affected side; 3, unable to bear weight on the affected side; and 4, no spontaneous locomotor activity or barrel rolling. If no deficit was observed 60 mins after beginning of occlusion period, the animal was removed from further study.

After 24 h reperfusion, the mice were sacrificed by rapid decapitation under deep anesthesia. Whole brain was rapidly removed for wet weight quantification, and then desiccated at 105°C for 48 h until the weight was constant for dry weight quantification. The water content and tissue swelling were calculated as follows based on [43]:

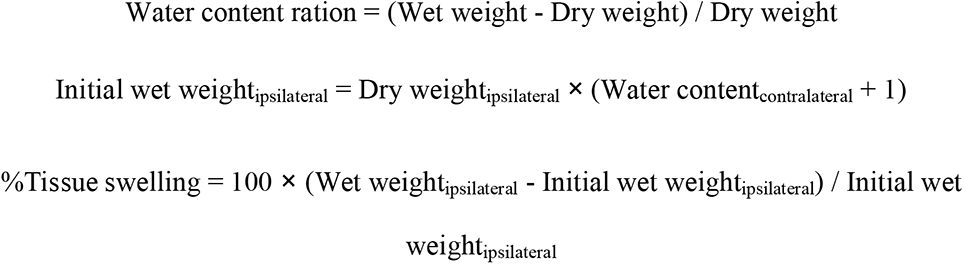

Brain infarct area was evaluated using 2, 3, 5-triphenyltetrazolium chloride (TTC) staining. Brains were sectioned into 2 mm-thick coronal slices and stained in 2% TTC at 37°C for 15 min in the dark and then photographed. The infarct tissue areas were measured using Image-Pro Plus. To account for edema, the infarcted area was estimated by subtracting the uninfarcted region in the ipsilateral hemisphere from the contralateral hemisphere, and the infarct volume was expressed as a percentage of the contralateral hemisphere.

### 4.9 ChIP and high-throughput sequencing

ChIP was performed as described previously [44]. Briefly, HT-22 cells were subjected to OGD treatment and crosslinked with 1% formaldehyde for 10 min at room temperature. Formaldehyde was quenched by 200 mM glycine and cells were rinsed twice with ice-cold PBS. Cells were transferred to 15 ml conical tubes and collected by centrifugation. Cells were lysed with lysis buffer A (50 mM HEPES-KOH (pH 7.5), 140 mM NaCl, 1 mM EDTA, 0.5% NP-40, 0.25% Triton X-100, 10% Glycerol and protease inhibitor cocktail (Roche)), incubated at 4°C for 10 min and collected by spinning at 1300 × *g* for 5 min at 4°C. Cells were then resuspended in lysis buffer B (10 mM Tris-Cl (pH 8), 200 mM NaCl, 1 mM EDTA, 0.5 mM EGTA and protease inhibitor cocktail), incubated at room temperature for 10 min. Nuclei were pelleted by spinning at 1300 × *g* for 5 min at 4°C. The pellet was suspended with lysis buffer B (10 mM Tris-Cl (pH 8), 100 mM NaCl, 1 mM EDTA, 0.5 mM EGTA, 0.1% Na-Deoxycholate, 0.5% N-lauroylsarcosine and protease inhibitor cocktail) and incubated for 15 min on ice. Chromatin shearing was conducted with cells on ice, using a microtip attached to Misonix 3000 sonicator. Sonicate 8-12 cycles of 30 s ON and 90 s OFF around 30-watt power-output. A final concentration of 1% Triton X-100 was added and gently mixed by pipetting. The chromatin solution was clarified by spinning at 20 000 *g* at 4°C for 30 min. Chromatin immunoprecipitation was performed with 50 ul Dynabeads protein G (Life technology) conjugated preliminary antibodies antibody overnight at 4°C. The immunoprecipitated material was washed five times with wash buffer (10 mM Tris-Cl (pH 8), 1 mM EDTA, 0.5% NP40, 0.5M LiCl, 0.5% Na-Deoxycholate) and once with TE buffer (PH 8.0), then, eluted by heating for 30 min at 65°C with elution buffer (50 mM Tris-Cl (pH 7.5), 10 mM EDTA, 1% sodium dodecyl sulphate). To reverse the crosslinks, samples were incubated at 65°C overnight, then the eluted was digested with a final concentration of 0.5 μg/ml RNasesA at 37°C, followed with a final concentrated of 0.5 μg/ml Proteinase at 55°C for 2 h. The immunoprecipitated DNA were then purified using the DNA clean and concentrator 5 column (Zymo Research). The ChIP DNA was used for qPCRs using indicated primers (Table S3) and data were plotted as the percentage of input. All experiments were performed three or more times, and representative results were shown in the figures. For ChIP-seq, 1 ng precipitated DNA or input was used to generate sequencing libraries using the Nextera XT DNA sample preparation Kit (Illumina). The resulting libraries were sequenced on Next-Seq (Illumina). Two biological replicates were performed, and combined reads were used for further analysis.

### 4.10 Bioinformatics analysis

For ChIP-Seq, reads were filtered if they had a mean Phred quality score of less than 20. They were aligned to the mm10 assembly using Bowtie v1.2 with parameters -v 2 -m 1 –best --strata. Reads that aligned to the same genomic coordinates were considered duplicates and removed using the MarkDuplicates tool in the Picard Tools suite v1.86. Fragment length estimates were obtained using Homer v4.3. Coverage tracks were generated using the genomecov tool in the BEDtools suite after extending the aligned reads to the estimated fragment lengths. Coverage tracks were normalized to coverage per 10 million mapped reads. Peaks were called using SICER v1.1 and the following parameters: species = mm10, redundancy threshold = 100, window size = 200, effective genome fraction = 0.77, gap size = 600, FDR = 0.000001. Peaks were labeled as “TSS” if they overlapped with the TSS of a gene in the GENCODE vM12 annotation, and they were labeled as “gene body” if they didn’t overlap with a TSS but did overlap with any other part of a gene body. They were labeled as “intergenic” if they didn’t overlap with either of the previous two.

For RNA-Seq, reads were filtered if they had a mean Phred quality score of less than 20. They were aligned to the mm10 assembly using STAR v2.6.0c. Gene counts were obtained using the featureCounts tool in the Subread package v1.5.1 with the GENCODE vM12 annotation. Differentially expressed genes (DEGs) were identified using DESeq2 with the following model: ∼ KnockDown + Treatment + KnockDown:Treatment. DEGs were required to have an FDR of less than 0.05 and a fold change of 1.5 or greater. “Down Reversed Genes” were those that had a significant negative fold change in the OGD *vs.* Control contrast and also a significant positive fold change in the interaction term contrast. The enrichment plot was generated using GSEA v.3.0.

### 4.11 Statistical analysis

All data were presented as means ± SEM. Statistical analyses were performed using student’s *t*-test, one-way analysis of variance (ANOVA) or two-way repeated-measures ANOVA followed by Bonferroni’s multiple comparison tests where appropriate. A value of *P* < 0.05 was considered as significant.

### 4.12 Data availability

All genomic datasets generated in this study (ChIP-Seq and RNA-seq) have been uploaded in the Gene Expression Omnibus (GEO) under accession number GSE248393.

## Acknowledgements

This section is not mandatory.

